# Comparative Analysis of Tick Microbiomes in Remnant and Reconstructed Prairie Ecosystems of Central Missouri

**DOI:** 10.1101/2025.07.06.663241

**Authors:** Mary A. York, Jaylon Vaugh, Deborah M. Anderson, Lyndon M. Coghill, Samniqueka J. Halsey

## Abstract

The ecological processes that shape tick-associated microbial communities are not fully understood. Ticks harbor diverse microbial communities that include both symbiotic and pathogenic bacteria, and these microbes can influence tick physiology, vector competence, and pathogen transmission. We evaluated bacterial microbiome diversity and composition in two medically important tick species, *Amblyomma americanum* and *Dermacentor variabilis*, collected from remnant and reconstructed tallgrass prairies in central Missouri between 2020 and 2023. Using 16S rRNA gene amplicon sequencing, we characterized microbial communities from 200 pooled tick samples and assessed patterns in alpha diversity, beta diversity, and differential abundance across tick species and prairie types. Microbiome diversity and composition differed strongly between tick species, with *D. variabilis* exhibiting higher alpha diversity than *A. americanum* and clear species-specific clustering in beta diversity analyses. In contrast, differences between remnant and reconstructed prairies were modest. Differential abundance testing revealed *Rickettsia* to be enriched in *A. americanum* and *Francisella* in *D. variabilis*, consistent across prairie types. Species-level screening further showed that *A. americanum* primarily harbored low virulent *Rickettsia*, whereas *D. variabilis* more frequently carried pathogenic *Rickettsia* species. Prairie restoration status did not significantly predict *Rickettsia* type, although a higher proportion of potentially pathogenic lineages was observed in reconstructed sites. These findings indicate that tick species identity is the primary determinant of microbiome structure in prairie ecosystems, with habitat context exerting secondary influences. Our results underscore the importance of incorporating microbiome ecology into medical entomology and suggest that land management practices may subtly shape vector–microbe interactions relevant to disease risk.

## Introduction

Tick-borne diseases are an expanding public health concern in the United States, driven by interacting changes in climate, land use, and host distributions(Eisen et al. 2017, Beard et al. 2019, Reaser et al. 2021). Across many regions, ticks serve as vectors for a diverse range of bacterial genera, including *Rickettsia* and *Francisella*, which can function as either symbionts or pathogens depending on the tick species and ecological context (Bonnet and Pollet 2021, Tonk-Rügen et al. 2023). Many of these bacteria are vertically transmitted and play integral roles in tick physiology, reproductive fitness, and vector competence, linking microbial ecology directly to disease transmission dynamics (Narasimhan and Fikrig 2015, Bonnet and Pollet 2021).

Beyond their role as disease vectors, ticks harbor complex microbial communities that are essential to their survival and development (Kolo and Raghavan 2023). Some microbial associates provide critical nutritional supplementation or developmental support, whereas others can disrupt microbial balance, alter immune function, and influence pathogen establishment and transmission (Adegoke et al. 2020). This dual role of symbionts and pathogens underscores the importance of studying tick microbiomes not only as reservoirs of infectious agents, but as structured ecological systems shaped by both host biology and environmental exposure (Wu-Chuang et al. 2021, Fountain-Jones et al. 2023).

Understanding tick-associated microbial communities is particularly important in regions where multiple vector species co-occur, as differences in host biology can lead to distinct microbiome composition and disease risk profiles (van Treuren et al. 2015, Fountain-Jones et al. 2023). In the central and southeastern United States, *Amblyomma americanum* and *Dermacentor variabilis* are among the most prevalent tick species, each known to harbor characteristic microbial assemblages (Trout Fryxell and DeBruyn 2016). Notably, *A. americanum* frequently carries *Rickettsia amblyommatis*, a highly prevalent low virulent species whose clinical significance in humans remains unresolved despite its widespread distribution (Eremeeva and Dasch 2015, Dahlgren et al. 2016). The dominance of species-specific microbial associates in these ticks highlights the need for focused comparative studies that link microbiome structure to host identity and ecological context.

Although host species identity plays a central role in shaping tick microbiomes, microbial community structure is not dictated by host biology alone (van Treuren et al. 2015, Brinkerhoff et al. 2020). A growing body of research demonstrates that ecological and environmental factors, including vegetation structure, host community composition, and microclimate, can influence microbial assemblages within ticks by altering exposure to environmental and host-associated microbial pools (Kueneman et al. 2021, Lejal et al. 2021). These effects are often subtle or indirect but may be especially relevant in human-modified landscapes where management practices and habitat restoration alter ecological interactions that shape vector and pathogen dynamics.

Despite increasing interest in the ecological drivers of tick microbiomes, relatively little is known about how prairie reconstruction, a widespread land management strategy in the central Midwest, influences microbial communities within ticks (Narasimhan, Swei, Abouneameh, Pal, Pedra, et al. 2021, Wimms et al. 2023). Most prior work has focused on forested or peri-urban systems, leaving restored grassland ecosystems underrepresented in studies of vector-associated microbiomes and disease risk (Brinkerhoff et al. 2020, Lejal et al. 2021). As prairie restoration continues to expand across the Midwest, understanding how these landscapes shape tick microbiomes represents a critical gap in our ability to predict and manage tick-borne disease risk in restored ecosystems (Reaser et al. 2021).

In this study, we compared the bacterial microbiomes of ticks collected from remnant and reconstructed prairie ecosystems to evaluate how restoration status shapes microbial community structure. Specifically, our objectives were to (1) compare alpha and beta diversity of tick microbiomes across prairie types, (2) assess whether microbial community composition differs by prairie type and tick species identity, and (3) identify bacterial taxa, including potential symbionts and pathogens, that are differentially abundant across environmental contexts. By integrating community diversity metrics, multivariate analyses, and differential abundance testing, this study provides new insight into how ecological restoration may influence host-associated microbial communities, with implications for understanding microbiome assembly processes and ecosystem health in restored landscapes.

## Methods

### Study Site

Tick sampling was conducted at two tallgrass prairie ecosystems in central Missouri: Tucker Prairie (remnant) and the Prairie Fork Conservation Area (PFCA) (reconstructed). These sites differ in ecological history, vegetation structure, and restoration management, making them ideal for investigating the effects of prairie restoration on tick-borne microbial communities. Tucker Prairie is a 146-acre remnant tallgrass prairie located in Callaway County, Missouri. Purchased by the University of Missouri (MU) in 1957, it remains the largest unplowed prairie in mid-Missouri. The site is actively managed by the MU Division of Biological Sciences in collaboration with the Missouri Department of Conservation (MDC). Tucker Prairie supports over 250 plant species across 70 families and 150 genera, with approximately 50% ground cover. Prairie Fork Conservation Area (PFCA) is a 911-acre complex of reconstructed prairie, savanna, and woodland habitats also located in Callaway County. Beginning in 2004, over 495 acres of prairie within PFCA were seeded with native grasses and forbs sourced from regional remnant prairies, including Tucker Prairie (Demand 2015).

### Questing Tick Collection

Ticks were collected monthly from May through August along five 100-meter transects established at each prairie site. Sampling was conducted using a standardized flagging method, in which a one-square-meter white corduroy cloth affixed to a wooden dowel was slowly dragged over herbaceous vegetation along both sides of each transect(Kjellander et al. 2021). This method covered a total sampling area of 1,000 m² per plot per visit. Every 15 meters, the cloth was examined for ticks, which were removed using masking tape and placed into plastic vials containing 70% ethanol. All collected ticks were first identified to species using standard taxonomic keys (Keirans and Durden 1998) and then sorted by species, life stage, and sex. Specimens were subsequently stored at −80°C until DNA extraction and microbiome analysis.

### DNA Extraction and Pooling

Ticks were initially pooled for DNA extraction based on site, plot, species, life stage/sex, and collection date (month and year). Each pool contained no more than five adult ticks or ten nymphs, depending on availability. DNA was extracted from these pools using the Quick**-**DNA/RNA™ Pathogen MiniPrep kit (Zymo Research) following the manufacturer’s protocol. After extraction, DNA concentration was measured using a Qubit Fluorometer (Invitrogen) with the Quant-iT™ High Sensitivity dsDNA Assay Kit. Extracted DNA from these initial pools was then aggregated into larger pools of up to six samples each, grouped by site, species, and life stage to ensure representation across environmental and biological conditions. This second-level pooling resulted in 436 aggregated DNA pools, which were submitted for 16S rRNA gene amplicon sequencing. Only DNA pools with concentrations greater than 2 ng/μL were retained for sequencing, yielding 200 high-quality samples for downstream microbiome analysis.

### 16S rRNA Sequencing and Bioinformatics

Bacterial community composition was assessed by sequencing the V4 hypervariable region of the 16S rRNA gene. Amplicons were generated using primers flanked by Illumina standard adapter sequences and unique dual indexes for each sample (Caporaso et al. 2011, Walters et al. 2011). Following PCR amplification, the amplicon libraries were quality-checked using an Advanced Analytical Fragment Analyzer, quantified, normalized, and pooled according to Illumina’s standard protocol for sequencing on the MiSeq platform. Raw sequencing reads were processed using a custom Nextflow v21.10.6 pipeline built around QIIME2 v2021+ (Di Tommaso et al. 2017). Primer removal was performed using Cutadapt (Martin 2011) and quality filtering, denoising, and identification of amplicon sequence variants (ASVs) were conducted using DADA2 both implemented within the QIIME2 platform (Callahan et al. 2016). Taxonomic assignment was performed using a Naive Bayes classifier trained on the SILVA 138 reference database (Pedregosa et al. 2011). To enable downstream taxon-level comparisons for differential abundance testing, ASV counts were aggregated to the genus level. Prior to diversity analysis, rarefaction was performed to standardize sequencing depth across samples at 15,400 ASV counts per sample. A phylogenetic tree of ASVs was also constructed using QIIME2 to support phylogeny-based diversity metrics and evaluate evolutionary relationships among bacterial taxa.

### Species-Specific Rickettsia Identification

We performed species-level Rickettsia identification via conventional PCR targeting the gltA gene using primers CS-78 (5′ -GCAAGTATCGGTGAGGATGTAAT - 3′) and CS-323 (5′ -GCTTCCTTAAAATTCAATAAATCA GGAT - 3′). Thermal cycling conditions were: an initial denaturation at 95°C for 5 minutes; followed by 40 cycles of 95°C for 15 seconds, 48°C for 30 seconds, and 72°C for 30 seconds; with a final extension at 72°C for 7 minutes. These conditions were based on previously published spotted fever group *Rickettsia* protocols (e.g., (Labruna et al. 2004)), with a slightly extended initial denaturation step to accommodate pooled tick DNA samples. PCR products were visualized on a 1.8% low-melt agarose gel run at 100V for 1 hour. Positive amplicons were excised and purified using the Wizard SV Gel and PCR Clean-Up System (Promega), then submitted for Sanger sequencing at the University of Missouri Genomics Technology Core. Resulting sequences were compared against the NCBI nucleotide database using BLASTn for species identification. For samples containing a single Rickettsia amplicon sequence variant (ASV) in 16S rRNA data, species identity determined by gltA sequencing was used to assign species identity to the corresponding ASV, which was subsequently applied as a sequence identifier across all samples containing that ASV.

### Statistical Analysis

We evaluated differences in microbiome diversity and composition among adult ticks across species (*Amblyomma americanum* and *Dermacentor variabilis*) and prairie site type (remnant vs. reconstructed).

#### Alpha and Beta Diversity

To evaluate how bacterial communities varied across tick species and prairie types, we analyzed both within-sample (alpha) and between-sample (beta) diversity. For alpha diversity, we calculated Observed ASVs, Chao1 richness, Shannon diversity, and the Inverse Simpson index (Chao 1984, Magurran 2021). We used Kruskal–Wallis tests to assess whether diversity differed between tick species or prairie sites, choosing this non-parametric approach because each factor had only two levels and data distributions were non-normal (Kruskal and Wallis 1952). To evaluate beta diversity, we transformed genus-level count data using a centered log-ratio (CLR) transformation after adding a pseudocount of one to account for zeros (Gloor et al. 2017). We then calculated Euclidean distances on the CLR-transformed matrix to quantify differences in community composition. Principal coordinates analysis (PCoA) was used to visualize clustering among samples, and we tested for significant differences using PERMANOVA with 999 permutations (Anderson 2001). Species, prairie site, and life stage were included as predictors in the model. To ensure that PERMANOVA results reflected true shifts in microbial composition and not differences in within-group variability, we used PERMDISP to test homogeneity of dispersion.

#### Differential Abundance

To identify bacterial taxa associated with tick species and prairie site, we conducted differential abundance analysis at the genus level using ANCOM-BC2, which accounts for compositionality and sampling bias in microbiome count data (Lin and Peddada 2020, 2024). Sequence counts were aggregated at the genus level prior to analysis, consistent with the taxonomic resolution of 16S rRNA amplicon data. ANCOM-BC2 models included tick species and prairie site as fixed effects, with life stage included as a covariate. P-values were adjusted for multiple testing using the Holm method (Holm 1979).

To stabilize variance estimates and avoid spurious significance associated with very small standard errors, a small constant corresponding to the 5th percentile of standard error values was added, following recommendations in the ANCOM-BC2 framework (Lin and Peddada 2024). Structural zero detection and asymptotic bounds were not applied, and statistical significance was assessed at α = 0.05. Although differential abundance testing was conducted across all detected genera, results are presented here only for *Rickettsia* and *Francisella*, which were the focal taxa for this study.

#### Pathogen-Associated Genera

We then tested whether the prevalence of pathogenic and endosymbiotic *Rickettsia* differed by site and tick species using chi-square tests of independence(McHugh 2013). For this analysis, we used presence-absence data from 105 tick pools, including 44 *Amblyomma americanum* and 61 *Dermacentor variabilis*, screened individually for *Rickettsia*. Each tick was classified as positive for either a pathogenic or endosymbiotic *Rickettsia* strain, based on species-level detection results.

## Results

From 2020-2023, we collected a total of 3,026 ticks from two prairies: Tucker Prairie (remnant, n = 1,097) and Prairie Fork Conservation Area (reconstructed, n = 1,929). Ticks belonged to two species: the lone star tick (*Amblyomma americanum*, n = 1,966; 65.0%) and the American dog tick (*Dermacentor variabilis*, n = 1,060; 35.0%). Ticks were nearly evenly distributed between life stages, with 1,561 nymphs (51.6%) and 1,465 adults (48.4%, Figure 1). Among adult ticks, 833 (56.9%) were female and 632 (43.1%) were male.

**Figure 1.**
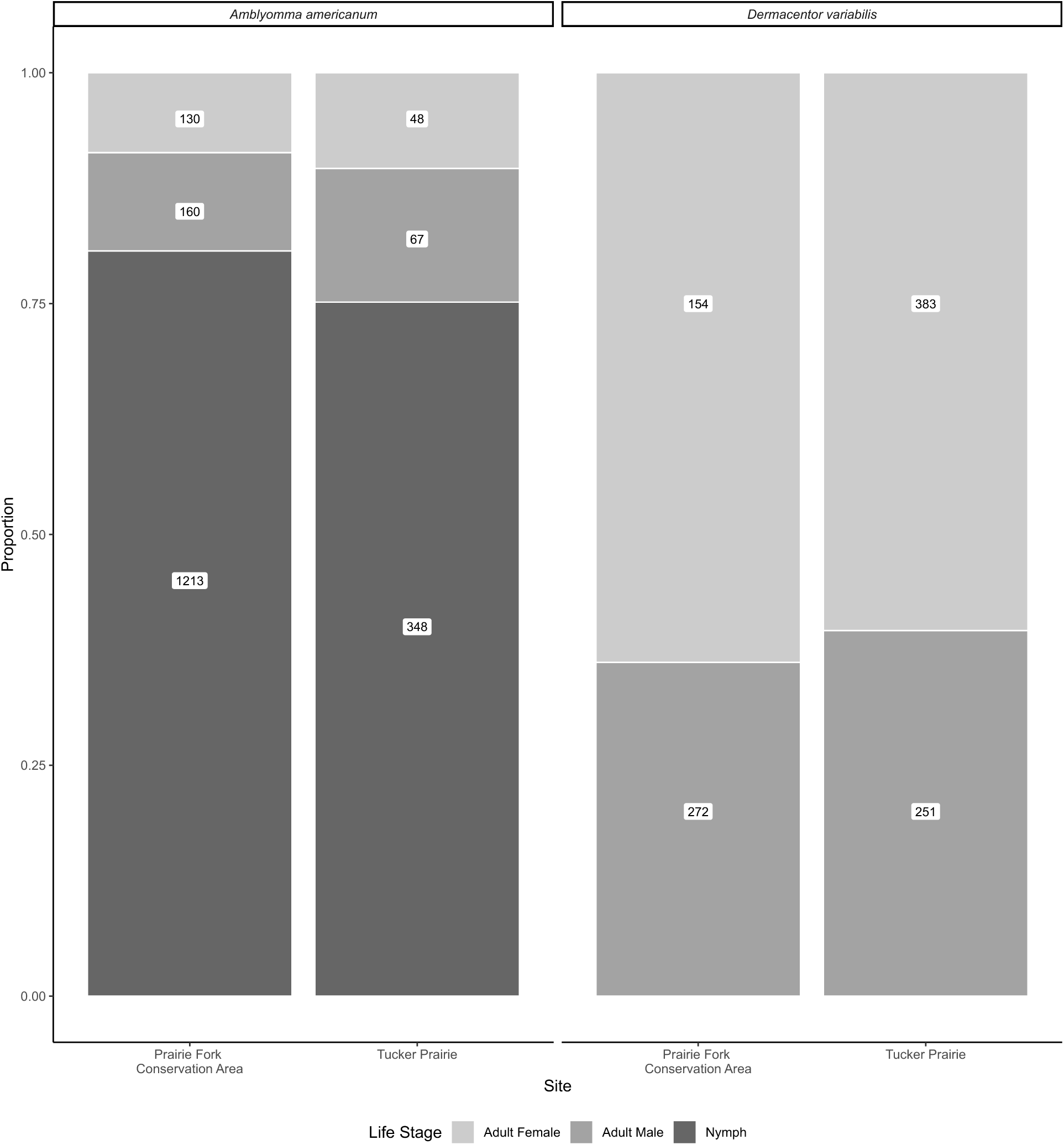
Proportional composition of tick life stages and sexes by species and prairie type from 2020 to 2023.Bars represent the relative proportions of Amblyomma americanum (Lonestar tick) and Dermacentor variabilis (American dog tick) collected between May and August, 2020–2023, from two prairie sites: Tucker Prairie (remnant) and Prairie Fork Conservation Area (reconstructed in Callaway County, Missouri. Numbers within each bar indicate the total number of ticks in each life stage and sex category.

A total of 436 tick pools were submitted for 16S rRNA gene amplicon sequencing. DNA extractions yielding <2 ng/μL, as quantified by Qubit fluorometry, were excluded from downstream analysis. All pools that met the DNA concentration threshold consisted of adult ticks, whereas pools containing only immature life stages (larvae or nymphs) consistently yielded insufficient DNA for sequencing. As a result, 200 adult tick pools were retained for sequencing. After demultiplexing and quality filtering, a total of 24,614,229 paired-end reads were generated across these samples, with per-sample read counts ranging from 2 to 1,300,937 (mean: 123,071; median: 59,588; SD: 180,149). All retained samples passed quality filtering, and sequencing depth was sufficient to characterize the bacterial communities associated with each tick pool. A total of 12,666 unique bacterial amplicon sequence variants (ASVs) were identified, spanning 38 phyla, 99 classes, and 1,080 genera. Several ASVs were classified within tick pathogen-associated genera of interest, including *Francisella spp.* (73.5% of samples), *Rickettsia spp.* (54.0%), *Ehrlichia spp.* (5.5%), *Borrelia spp.* (1.5%), *Anaplasma spp.* (4.5%), and *Bartonella spp.* (1.5%).

### Alpha diversity

There were no significant differences in alpha diversity between sites for any metric (Observed: χ² = 0.007, *p* = 0.93; Chao1: χ² = 0.007, *p* = 0.93; Shannon: χ² = 0.007, *p* = 0.93; Inverse Simpson: χ² = 0.074, *p* = 0.79, Figure 2). In contrast, Shannon and Inverse Simpson indices differed significantly by tick species, with higher diversity observed in *Dermacentor variabilis* compared to *Amblyomma americanum* (Shannon: χ² = 7.92, *p* = 0.0049; Inverse Simpson: χ² = 14.15, *p* < 0.001). No significant differences by species were observed for richness-based metrics (Observed: χ² = 0.50, *p* = 0.48; Chao1: χ² = 0.50, *p* = 0.48).

**Figure 2.**
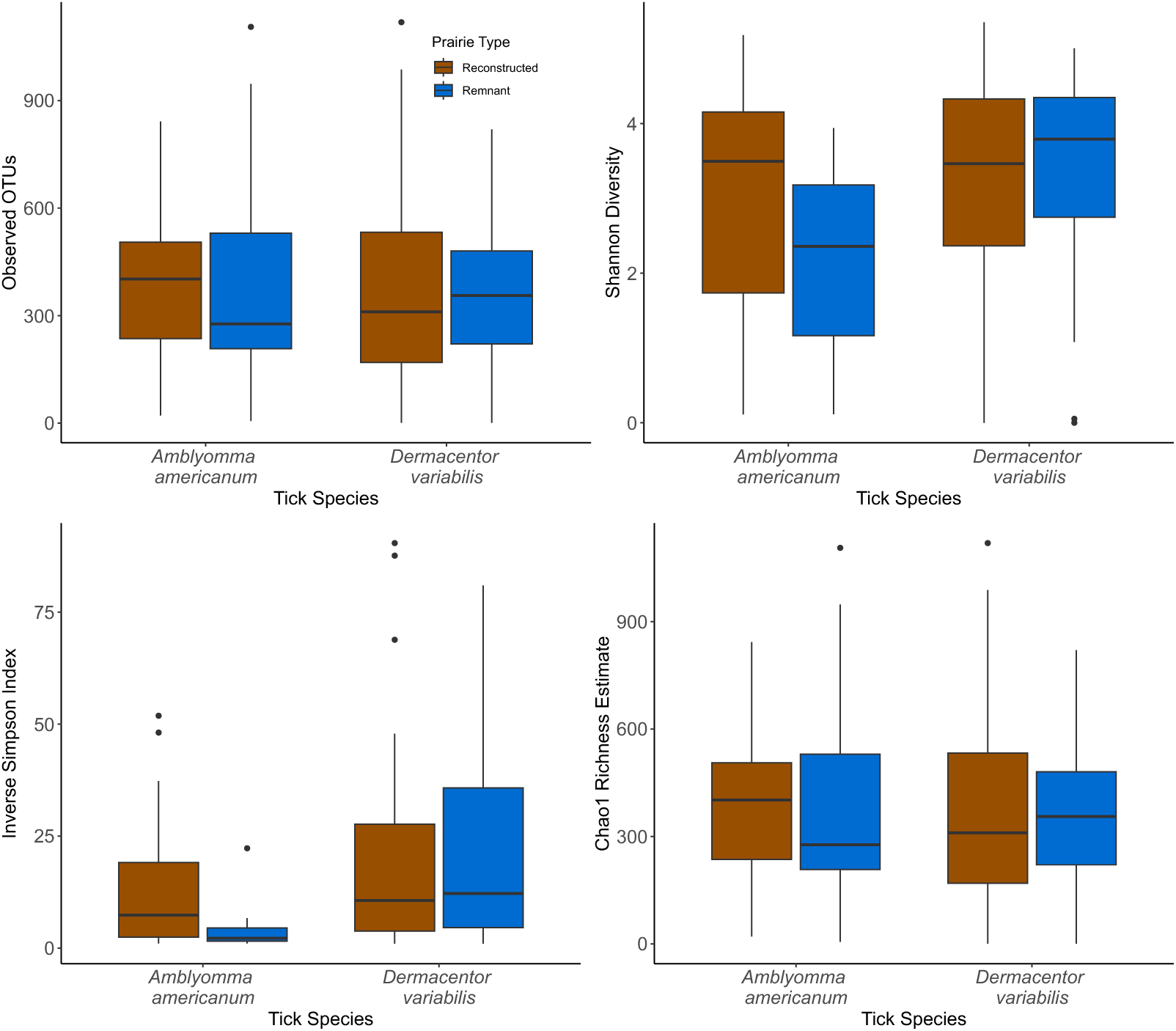
Alpha diversity of tick-associated bacterial communities by species and prairie type. Boxplots show four alpha diversity metrics—Observed ASVs, Chao1 richness estimate, Shannon diversity, and Inverse Simpson index stratified by tick species and prairie site (Reconstructed = Prairie Fork, Remnant = Tucker Prairie)..

### Beta Diversity

There were significant differences in the microbiome composition between remnant and reconstructed prairie (F_(1, 139)_ =2.206, *p* = 0.026) and *A. americanum* and *D. variabilis* (F_(1, 139)_ =35.709, *p* = 0.001). We found clustering in the bacterial microbiome between *A. americanum* and *D. variabilis* at the genus level (Figure 3), indicating that different tick species have distinct microbiomes. Community composition differed significantly by tick species (R² = 0.040, F_(1, 139)_ = 8.23, *p* = 0.001) and to a lesser extent by prairie site (R² = 0.010, F_(1, 199)_ = 2.06, *p* = 0.024), based on PERMANOVA of CLR-transformed genus-level bacterial communities. Tick life stage was not a significant predictor of microbial composition (R² = 0.005, F_(1, 199)_ = 1.02, *p* = 0.598). A combined model including tick species, site, and their interaction explained 5.7% of the total variation (F_(1, 199)_ = 3.94, p = 0.001), suggesting that microbiome structure is influenced by both host identity and environmental context. A PERMDISP test for homogeneity of group dispersions showed no significant difference in dispersion between tick species (F = 0.68, *p* = 0.41), supporting the interpretation that observed PERMANOVA differences reflect true shifts in community structure rather than within-group variability.

**Figure 3.**
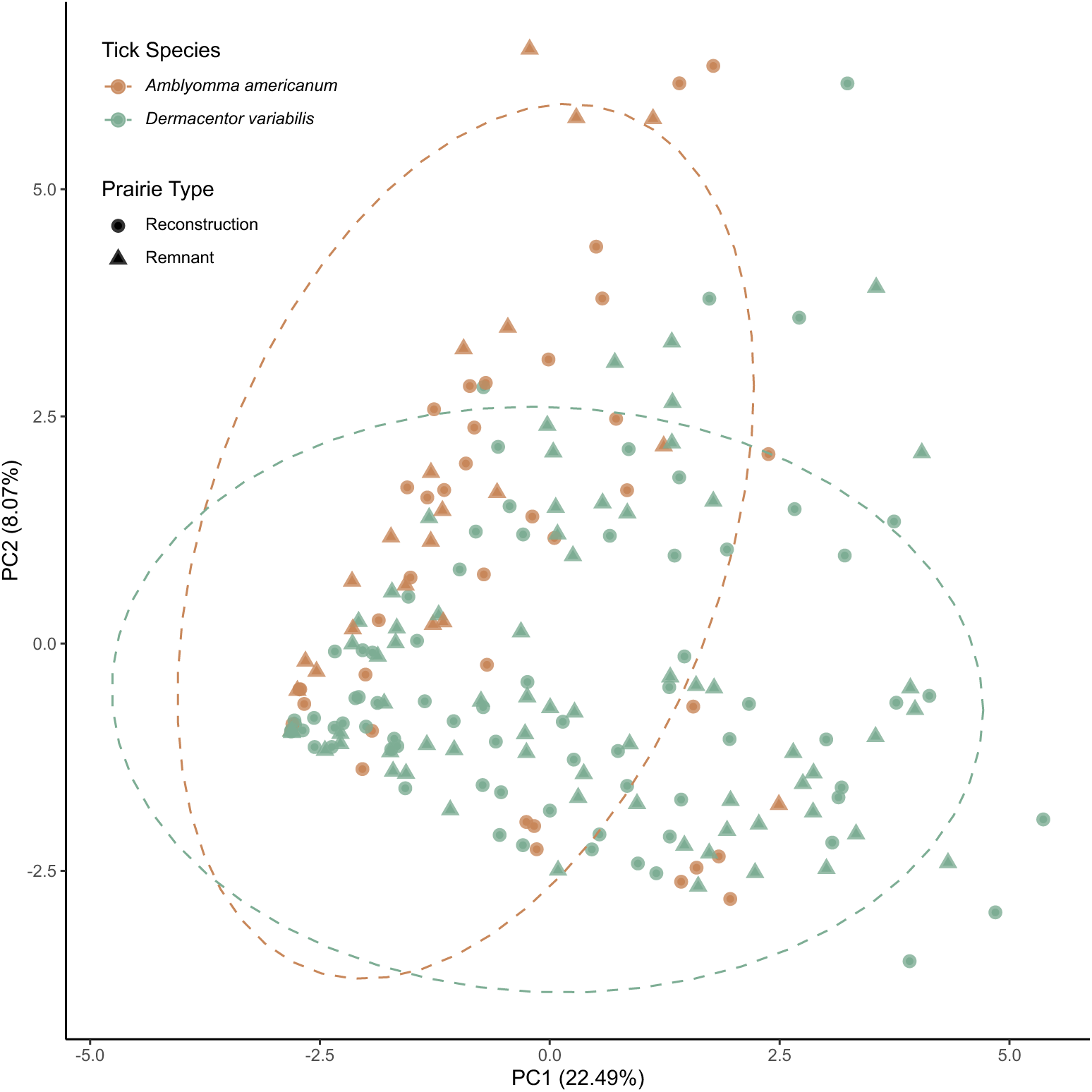
Principal component analysis (PCA) of genus-level CLR-transformed bacterial communities from ticks. Ordination is based on Euclidean distances calculated from centered log-ratio (CLR) transformed genus-level abundances. Each point represents a tick sample, colored by species (Amblyomma americanum [AA] and Dermacentor variabilis [DV]) and shaped by prairie site (Reconstructed = Prairie Fork, Remnant = Tucker Prairie). Dashed ellipses represent 95% confidence intervals around species centroids.

Differential abundance testing using ANCOM-BC2 revealed significant shifts in the relative abundance of *Rickettsia* and *Francisella* between tick species with higher abundance of *Rickettsia* in *Amblyomma americanum* compared to *Dermacentor variabilis* (*W* = −4.05, *p* < 0.001, Figure 4A). Conversely, *Francisella* was significantly more abundant in *D. variabilis* than in *A. americanum* (*W* = 4.18, *p* < 0.001). Furthermore, for *Rickettsia,* there was no significant difference in abundance between prairie sites (*W* = 0.56, *p* = 0.577), whereas for *Francisella*, a marginal, non-significant trend was observed (*W* = –1.67, *p* = 0.097), likely driven by *D. variabilis (*Figure 5). Prairie site was not a significant predictor of abundance for either genus, and life stage effects were not supported after adjustment for multiple comparisons.

**Figure 4.**
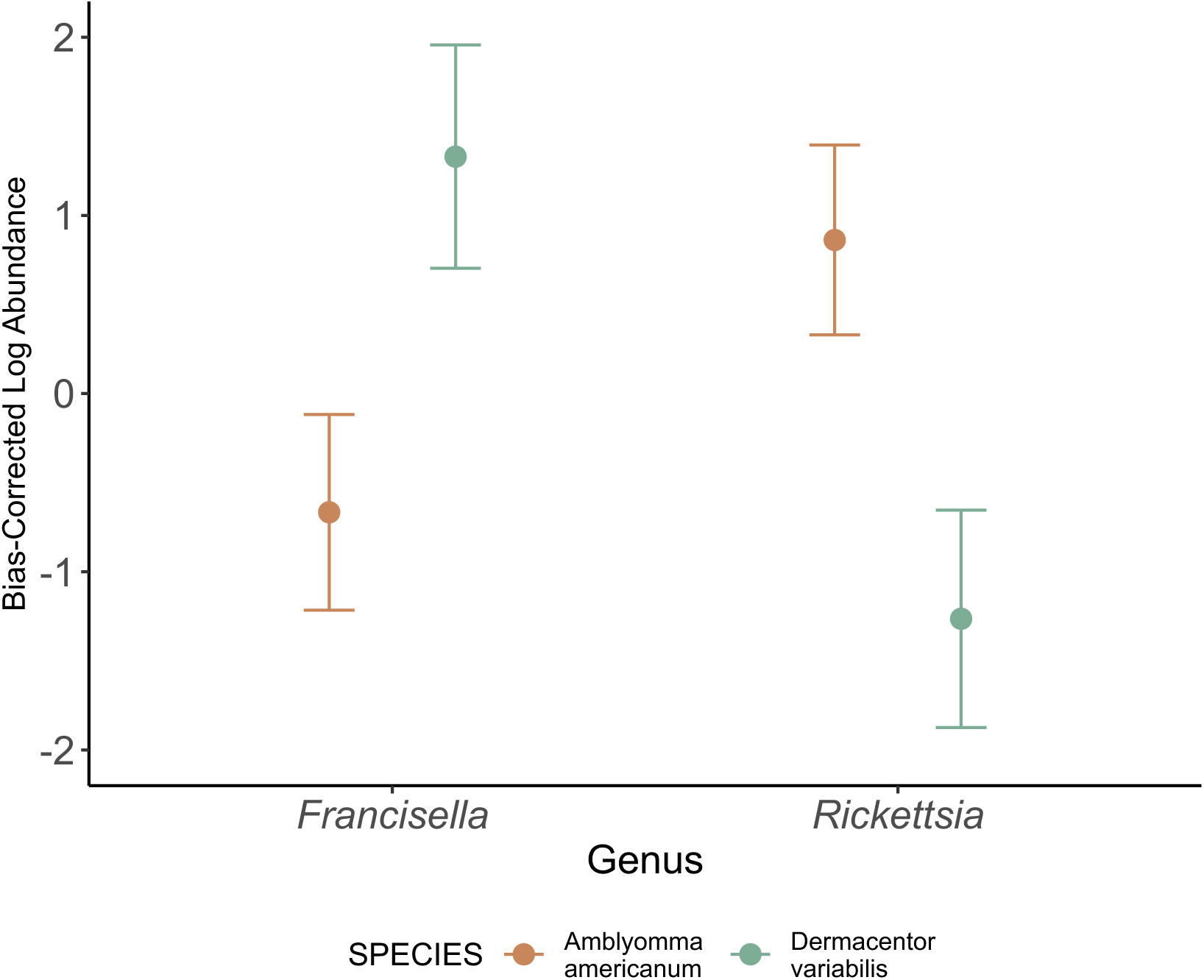
Mean Bray-Curtis Log log-transformed abundances (±95% confidence intervals) of differentially abundant genera between *Amblyomma americanum* and *Dermacentor variabilis*.

**Figure 5.**
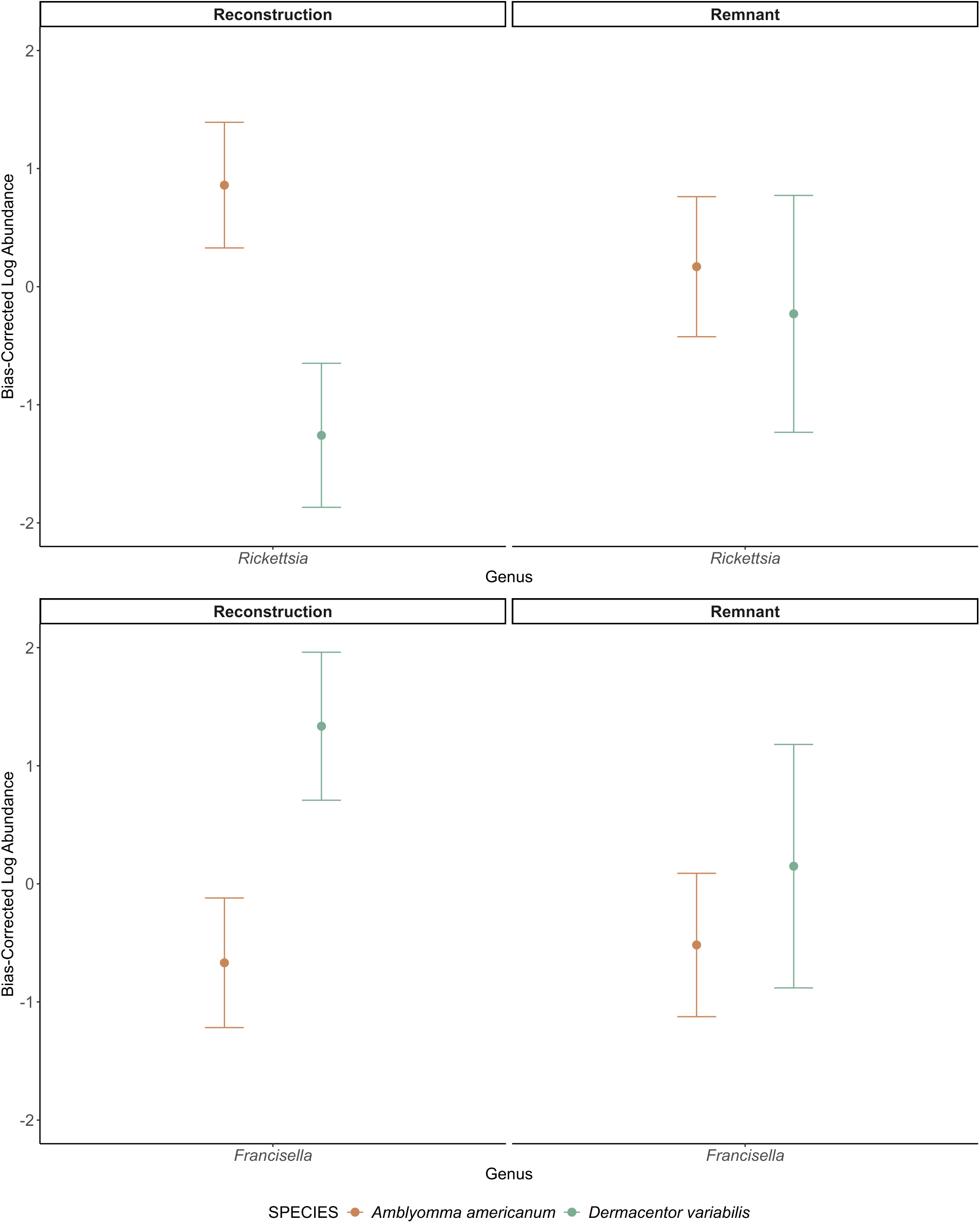
Mean Bray-Curtis Log log-transformed abundances (±95% confidence intervals) of differentially abundant genera (*Rickettsia spp*.) between Amblyomma americanum and Dermacentor variabilis.

### Pathogenic and low virulent Rickettsia prevalence

Of the 200 pools tested, 108 (54%) were positive for *Rickettsia*, and species-level identification was achieved for 105 of these, encompassing four distinct *Rickettsia* species: *R. amblyommatis*, *R. rickettsii*, *R. montanensis*, and *R. adanae*. Co-infection with multiple *Rickettsia* species was rare, occurring in only 5 of the 200 samples (2.5%). These included three *Dermacentor variabilis* and two *Amblyomma americanum* ticks, with three collected from the reconstructed prairie and two from the remnant site. No consistent pattern of co-occurrence was observed among the species-level combinations.

We found strong species-specific patterns in *Rickettsia* infection. Among the 44 *Amblyomma americanum* individuals, pathogenic *Rickettsia* were detected in only 5 ticks (11.4%), while less virulent species were detected in 41 (93.2%). In contrast, of the 61 *Dermacentor variabilis* ticks, 46 (75.4%) were positive for pathogenic *Rickettsia* and 15 (24.6%) carried low virulent. A chi-square test confirmed that pathogen type was significantly associated with tick species identity (χ² = 45.45, *p* < 0.001). When stratified by site, we observed that 55.6% of *Rickettsia*-positive ticks in Tucker Prairie carried pathogenic strains compared to 41.2% in Prairie Fork, but this difference was not statistically significant (χ² = 1.30, *p* = 0.254). These patterns suggest that *A. americanum* is primarily associated with low virulent *Rickettsia*, whereas *D. variabilis* more frequently carries pathogenic strains.

## Discussion

Tick species identity, rather than prairie restoration status, was the primary determinant of microbiome structure in *Amblyomma americanum* and *Dermacentor variabilis*. Consistent with prior studies, microbiome diversity and composition differed strongly between species (Brinkerhoff et al. 2020, Bonnet and Pollet 2021, Fountain-Jones et al. 2023), with *D. variabilis* exhibiting higher alpha diversity and distinct community structure compared to *A. americanum*, while site-level differences were comparatively modest. These species-specific patterns extended to dominant bacterial taxa, with *Francisella* enriched in *D. variabilis* and *Rickettsia* dominating *A. americanum* (Bonnet and Pollet 2021, Hensley et al. 2021). Prairie restoration status did not significantly predict *Rickettsia* type, and sex-associated differences in *Rickettsia* or *Francisella* abundance were not supported when life stage was included as a covariate. Together, these results reinforce the conclusion that host species identity outweighs environmental context in structuring tick microbiomes in prairie ecosystems (van Treuren et al. 2015, Brinkerhoff et al. 2020).

Although the effect was not statistically significant, we observed a trend toward a greater representation of potentially pathogenic *Rickettsia* lineages in reconstructed sites. This pattern is consistent with growing evidence that habitat structure and restoration can subtly modulate host–microbe interactions without necessarily driving large shifts in overall microbial diversity or composition (Kueneman et al. 2021, Lejal et al. 2021)

The higher alpha diversity observed in *D. variabilis* likely reflects species-specific differences in host use, environmental exposure, and immune-mediated microbial filtering (Bonnet and Pollet 2021, Fountain-Jones et al. 2023). Previous studies have shown that tick species identity and developmental stage consistently shape microbiome diversity, often more strongly than local environmental conditions (Brinkerhoff et al. 2020, Narasimhan, Swei, Abouneameh, Pal, Fikrig, et al. 2021). These patterns are thought to arise because species identity acts as a biological filter, with innate immune responses and vertically transmitted symbionts constraining microbial establishment and persistence within ticks (Hawlena et al. 2013, Bonnet and Pollet 2021).

Although tick species identity dominated microbiome structure, we detected modest but consistent site-level differences in beta diversity between remnant and reconstructed prairies. These differences likely reflect variation in microclimate, vegetation structure, and host community composition across prairie types, which can influence tick contact with environmental and host-associated microbial pools (Kueneman et al. 2021, Lejal et al. 2021). Vegetation heterogeneity and restoration status may therefore modulate tick microbiomes indirectly(Lejal et al. 2021), shaping community composition without overriding the strong species-specific constraints imposed by the tick host (Bonnet and Pollet 2021).

Species-specific dominance of obligate low virulent species further explains why host identity outweighed habitat effects in structuring tick microbiomes (Narasimhan, Swei, Abouneameh, Pal, Fikrig, et al. 2021). *Francisella* was enriched in *Dermacentor variabilis*, consistent with its established role as a dominant endosymbiont in *Dermacentor* spp.(Bonnet and Pollet 2021). These symbionts are thought to contribute to essential nutrient provisioning, particularly B vitamins, and to influence colonization resistance against other microbes (Tonk-Rügen et al. 2023). In contrast, *Amblyomma americanum* microbiomes were dominated by *Rickettsia*, primarily *R. amblyommatis*, a vertically transmitted symbiont associated with nutritional supplementation and potential reproductive benefits (Brinkerhoff et al. 2020). The stability of *Rickettsia* and *Francisella* abundance across prairie types indicates that these symbiotic associations are largely conserved across environmental contexts (van Treuren et al. 2015, Hensley et al. 2021). This pattern supports the broader hypothesis that obligate blood-feeding arthropods maintain stable, host-specific symbiont assemblages that are buffered against local habitat variation (Narasimhan, Swei, Abouneameh, Pal, Fikrig, et al. 2021).

Species-specific patterns in *Rickettsia* infection have clear implications for disease risk assessment (Eremeeva and Dasch 2015). *Dermacentor variabilis* is a recognized vector of pathogenic *Rickettsia* species, including *R. rickettsii* and *R. montanensis* (Dahlgren et al. 2016), whereas *Amblyomma americanum* more commonly harbors *R. amblyommatis*, a widespread symbiont of uncertain clinical relevance (Trout Fryxell and DeBruyn 2016). Although prairie restoration status was not significantly associated with *Rickettsia* type, we observed a higher proportion of potentially pathogenic lineages in reconstructed sites. This trend suggests that ecological restoration may subtly influence microbe-vector interactions through environmental filtering, host community turnover, or habitat-specific factors, even when overall patterns are dominated by host species identity (Kueneman et al. 2021, Lejal et al. 2021)

These findings add to growing evidence that ecological restoration can influence tick–microbiome interactions, even when host species identity remains the dominant determinant of microbial community structure (Brinkerhoff et al. 2020, Bonnet and Pollet 2021, Lejal et al. 2021). While tick species identity remains the strongest predictor of microbiome composition, modest site-level effects indicate that local habitat conditions may exert secondary influences. Vegetation structure, soil microbiota, and host diversity are plausible mediators of these patterns (Kueneman et al. 2021). This work reinforces the need for an integrated approach to vector management that considers microbial ecology alongside landscape restoration. Recognizing the interactions among ticks, microbes, and their habitats may enhance predictions of pathogen emergence and inform ecologically grounded public health strategies (Reaser et al. 2021).

Several aspects of the study design constrain the resolution at which tick-microbiome interactions could be evaluated. Pooling of samples limited inference about individual-level variation and reduced sensitivity to rare microbial taxa, while exclusion of larval stages precluded direct assessment of early-life symbiont acquisition and vertical transmission dynamics (Zhou et al. 2023, Gaire et al. 2025). In addition, reliance on 16S rRNA amplicon sequencing restricted strain-level resolution and remains susceptible to biases associated with DNA extraction efficiency, primer amplification, and taxonomic assignment. These limitations are inherent to many tick microbiome studies and underscore the need to interpret community-level patterns as indicators of dominant ecological processes rather than fine-scale microbial interactions (Bonnet and Pollet 2021)

Future work should build on these findings by increasing resolution across both biological and temporal scales. Individual-level sequencing that includes larval stages would allow direct assessment of microbiome assembly across development and clarify the role of vertical transmission in structuring tick-associated microbial communities. Coupling microbiome analyses with host blood meal identification and small mammal surveys would further disentangle vertebrate contributions to microbial acquisition and persistence. Beyond taxonomic profiling, functional metagenomic and transcriptomic approaches could reveal microbial interactions, metabolic roles, and activity in situ. Finally, longitudinal sampling across seasons and management regimes would help resolve how disturbance processes such as fire influence microbial community dynamics over time (Lejal et al. 2021, Tonk-Rügen et al. 2023).

## Acknowledgements

We would like to thank the numerous technician over the years for their help in sample collection, processing and DNA extraction. We thank Prairie Fork Conservation Area and Tucker Prairie for access to conduct this research in their prairies.

